# Tool use by four species of Indo-Pacific sea urchins

**DOI:** 10.1101/347914

**Authors:** Glyn Barrett, Dominic Revell, Lucy Harding, Ian Mills, Axelle Jorcin, Klaus M. Stiefel

## Abstract

We compared the covering behavior of four sea urchin species, *Tripneustes gratilla, Pseudoboletia maculata, Toxopneutes pileolus,* and *Salmacis sphaeroides* found in the waters of Malapascua Island, Cebu Province and Bolinao, Panagsinan Province, Philippines. Specifically, we measured the amount and type of covering material on each urchin, and, in several cases, the recovery of debris cover after stripping the animal of its cover.

We found that *Tripneustes gratilla* and *Salmacis sphaeroides* have a higher preference for plant material, especially sea-grass, compared to *Pseudoboletia maculata* and *Toxopneutes pileolus*, which prefer to cover themselves with coral rubble and other calcified material. Only for *Toxopneutes pileolus* did we find a decrease in cover with depth, confirming previous work that the covering behavior serves UV protection. We found no dependence of particle size on either species or urchin size, but we observed that larger urchins carried more and heavier debris. We observed a transport mechanism of debris onto the echinoid body surface utilizing a combination of tube feet and spines. The transport speed of individual debris items varied between species.

We compare our results to previous studies of urchin covering behavior, comment on the phylogeny of urchin covering behavior and discuss the interpretation of this behavior as animal tool use.

## Introduction

Several species of sea urchins cover their exposed body surfaces with debris from their environment (Agatsuma, 2001; Dumont *et al.*, 2007; Zhao, Ji, *et al.*, 2014; Wei *et al.*, 2016; Zhao, Bao and Chang, 2016; Ziegenhorn, 2016). The debris can include coral rubble, mollusk shells, sea grass and human rubbish. The urchins actively grab these debris elements with their tube feet and spines and transport them onto their upward facing body surface. Suggested reasons for this behavior are UV protection (demonstrated for *Tripneustes gratilla,* (Belleza, Samuel and Jr, 2012; Ziegenhorn, 2016)), olfactory camouflage (suggested for covering species living in dark deep-sea environments, (David, Magniez and Villier, 1999; Brothers *et al.*, 2016)) and use as a ballast for weighing down of the animal in the face of currents or surge (shown for *Strongylocentrotus droebachiensis,* (Dumont *et al.*, 2007)) as well as direct anti-predatory function (Zhao, Ji, *et al.*, 2014)(Agatsuma, 2001). Furthermore, since the covering material is often edible for the urchins (sea grass, coralline algae), covering has been suggested as a food storage behavior (Dix, 1970; Douglas, 1976). It has been shown for UV radiation and wave action (Dumont *et al.*, 2007; Belleza, Samuel and Jr, 2012), that urchins increase their coverage if faced with these stressors. Different species of urchins have been shown to be selective for the covering material used (Amato *et al.*, 2008).

Here we study and compare the covering behavior of four species, *Tripneustes gratilla, Pseudoboletia maculata, Salmacis sphaeroides* and *Toxopneutes pileolus*, species found near reefs at shallow depths in the tropical Indo-Pacific, specifically the Philippines. These species partially overlap in their habitat use, hence they have access to the same type of covering material. This makes differences in their covering behavior intriguing. We combine field observations of debris amount and type with video recordings of coverage material handling both in the field and in aquaria.

Given the active choice of coverage material and the situation-dependent loading of material, as well as the coordinated lifting of coverage material, we propose in the discussion that this behavior constitutes animal tool use.

## Methods

Echinoids were sampled on scuba at depths between 2 and 16 meters around Malapascua Island, Cebu Province, and Bolinao, Pangasinan Province, Philippines. All four investigated species are moderately common at the sampling sites. Of the 6 sites which we sampled, one site each was populated by 4, 3 and 1 species of the collector urchins we investigated, and three sites (including the sampling site in Bolinao) by 2 species. In the field, each urchin was photographed, measured (diameter), and carefully stripped of its attached debris. The debris was placed in a plastic bag. The “naked” urchin was then observed and in many cases filmed for up to 10 minutes, recording its covering behavior. If an urchin failed to re-cover itself we would place debris comparable to the removed debris on the animal to avoid stress to the urchin. Post dive, collected debris fragments were laid onto white slates with a ruler for scale and photographed from above. Individual pieces were counted, categorized, and surface area was determined using ImageJ (Schneider et al., 2012) (Fig.1). Categories were coral rubble, mollusk shells, tunicate, marine plants and algae, land plants, human refuse. Debris was sun-dried (24 hours) and weighted.

**Figure 1:**
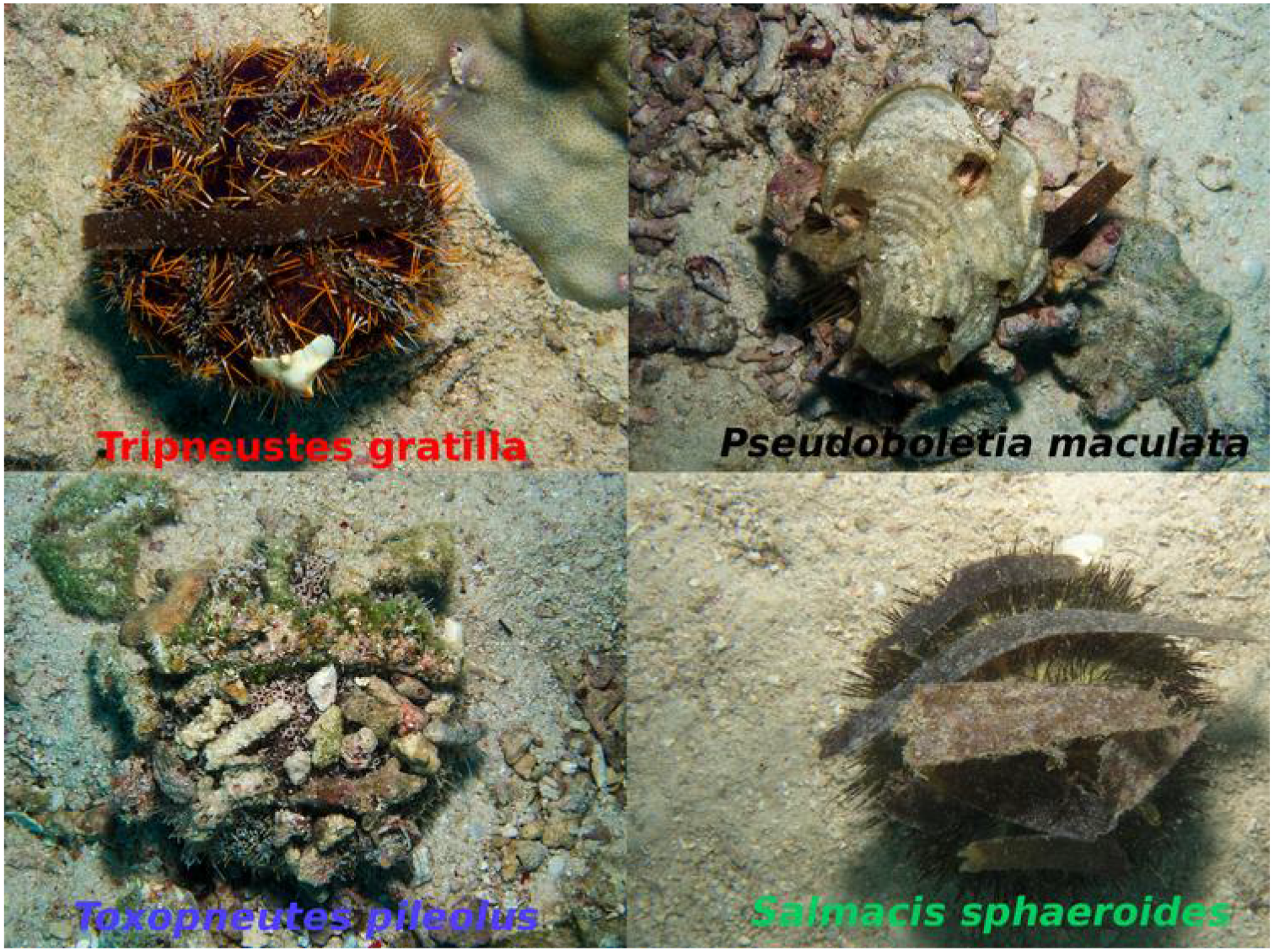
Examples of urchins covered in debris, photographed in their natural environment: *Toxopneutes pileolus, Tripneustes gratilla, Salmacis sphaeroides, Pseudoboletia maculata.*

To determine the speed of re-covering, we traced the position of selected pieces of debris during the urchin’s efforts to cover its body surface with debris. Footage was either shot in the field, by a diver with a hand-held camera, or in tanks, with a camera placed on bars above the tank.The video analysis was done with the Tracker software (Douglas Brown, physlets.org/tracker). At intervals of 10 seconds, the position of the urchin’s pole and the pole-most point of the fragment were manually indicated, and the resulting positions were used for the analysis of fragment movement speed. At each time step, the scale bar was also adjusted if necessary (if the distance from the camera to the urchin slightly changed in the case of hand-held footage, Fig. 6A). The radial position of the fragments determined in this way is an underestimation of the actual distance traveled, since the urchin bodies are compressed spheres. The measured speeds and positions are useful for comparative purposes, though.

Six urchins each of the species *Tripneustes gratilla* and *Salmacis sphaeroides* were collected in Bolinao and maintained in aquaria at the Marine Lab of the University of the Philippines. The urchins were fed with sea-grass and given sea-grass and coral rubble as coverage materials. Close-up views of the covering behavior were filmed in the tanks.

Some of the video footage analyzed for this study can be seen at https://www.youtube.com/watch?v=GZw0mKy_un8&t=4s

## Results

We sampled a total of 49 urchins, *T. gratilla* (n=11), *P. maculata* (n=10), *S. sphaeroides* (n=12) and *T. pileolus* (n=16). We compared debris type preference between species of collector urchins and found significant differences in coverage material most importantly between total organic (*e.g.* sea-grass, rodoliths, tunicata) (ANOVA, F (3, 45) = 7.198, *p* < 0.001) and total inorganic *(e.g.* coral, shell, rock) (ANOVA, F (3, 45) = 8.654, p < 0.001) materials (Fig. 2). *S. sphaeroides* preferred seagrass as a covering material whilst *T. pileolus* preferred coral fragments (Tukeys HSD posthoc).

**Figure 2:**
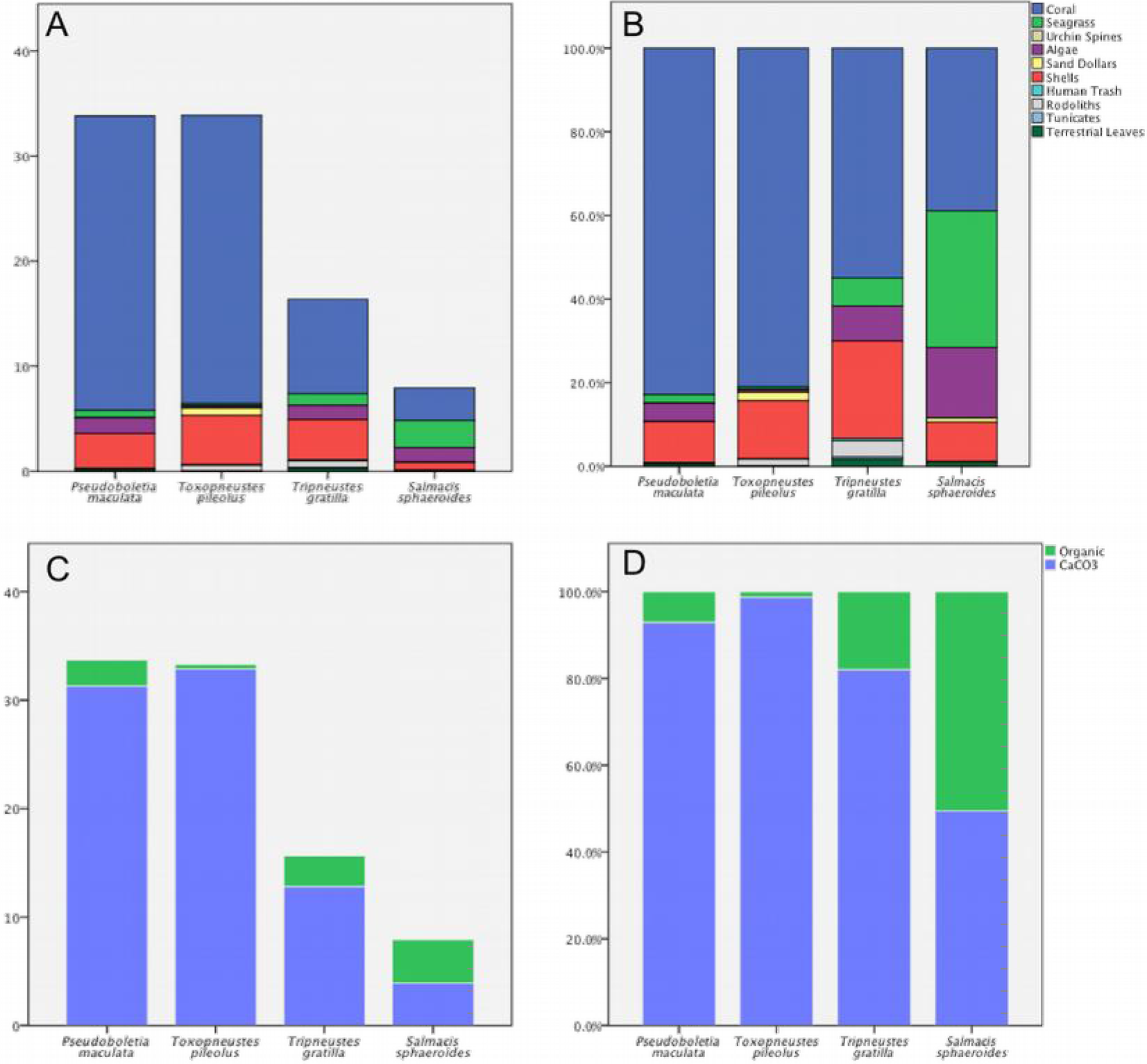
Different species of collector urchins prefer different types of debris to cover themselves. Species vs. debris type. Y-axis showing the absolute debris item numbers (A) the percentage of debris items (B), the absolute number of fragments sorted between calcified skeletons and organic material (C), and sorted fragments, in percent (D).

Linear regression was calculated to predict total cover area (cm^2^) based on depth (m) (Fig. 3). A significant regression equation was found for *T. pileolus* (F(1,14) = 7.443, *p* = 0.016), with an R^2^ of 0.347. *T. pileolus* predicted total cover area is equal to 248.4 - 10.1 (depth) cm^2^ when depth is measured in m. Urchin cover area decreased by 10.1 cm^2^ for each meter of depth. Depth was not a significant factor in influencing total cover area for *T. gratilla*, *P. maculata* or *S. sphaeroides*.

**Figure 3:**
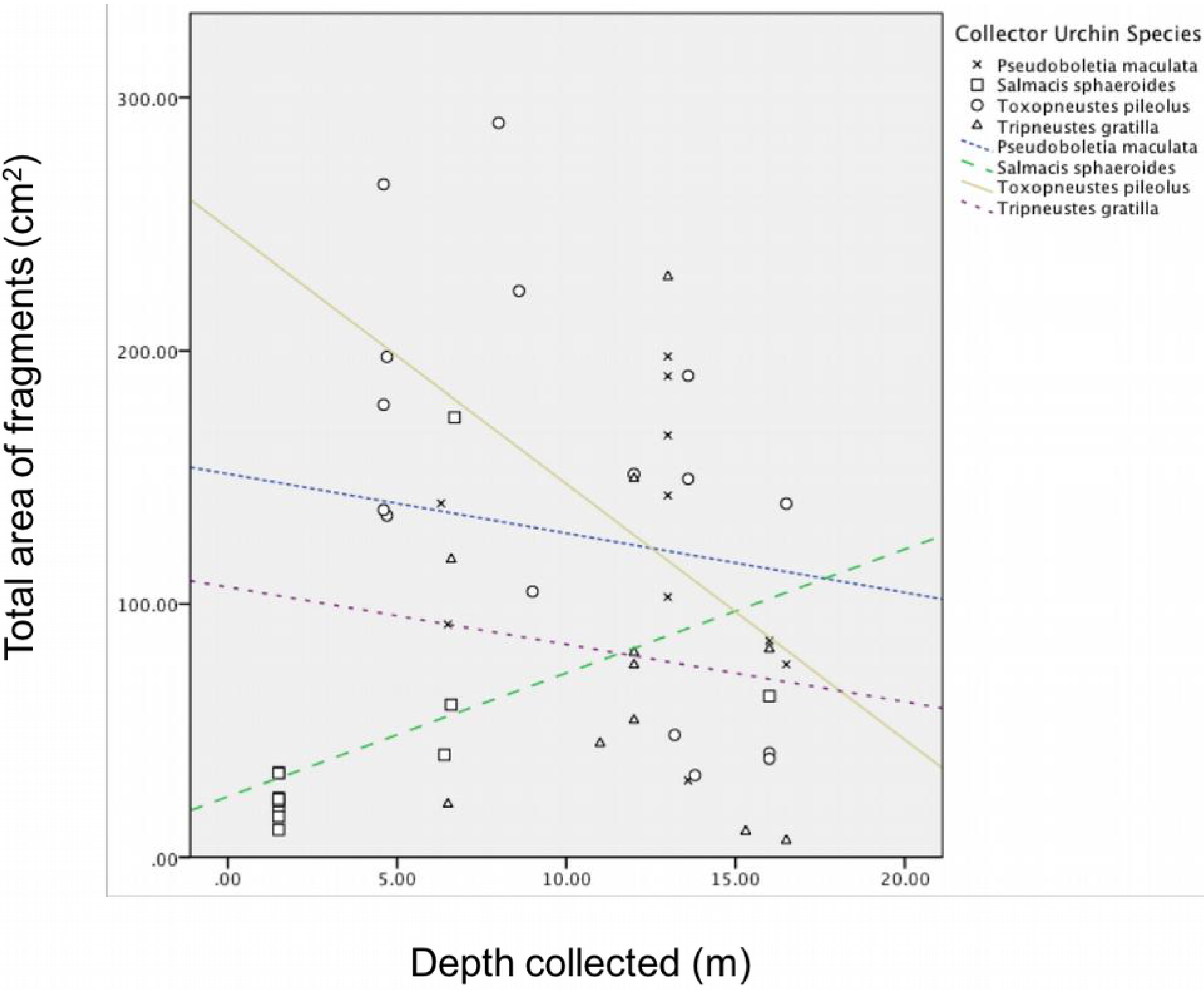
Depth influences the amount of debris on collector urchins. Fragment area for *Tripneustes gratilla, Pseudoboletia maculata, Salmacis sphaeroides* and *Toxopneutes pileolus.*

We found significant differences in size, as defined by equatorial diameter, of species of urchin (ANOVA, F(3, 45) = 211.722, *p* < 0.001) with *S. sphaeroides, P. maculata, T. gratilla* and *T. pileolus* at 6.3, 9.6, 9.7 and 14.8 cm, respectively (Fig. 4). Interestingly, the global mean size of fragments (5.6 ± 0.68 cm^2^) was independent of urchin species and size of urchin (ANOVA, F(1,48) = 0.7, p = 0.407) (Fig.5). Linear regression was calculated to predict both total number (n) and total weight (g) of fragments based on diameter of urchin (cm). Significant regression equations were found for both total fragment number (F(1,47) = 10.094, p = 0.003), with an R^2^ of 0.177 and total weight of fragments (F(1,47) with an R^2^ of 0.078. Predicted total fragment number is equal to 8.818 + 0.071 (diameter) n when diameter is measured in cm. Predicted total fragment weight is equal to 9.676 + 0.011 (diameter) g when diameter is measured in cm. Total fragment number and total fragment weight increased, respectively, by 0.071 and 0.011 g for each increase of 1 cm in diameter.

**Figure 4:**
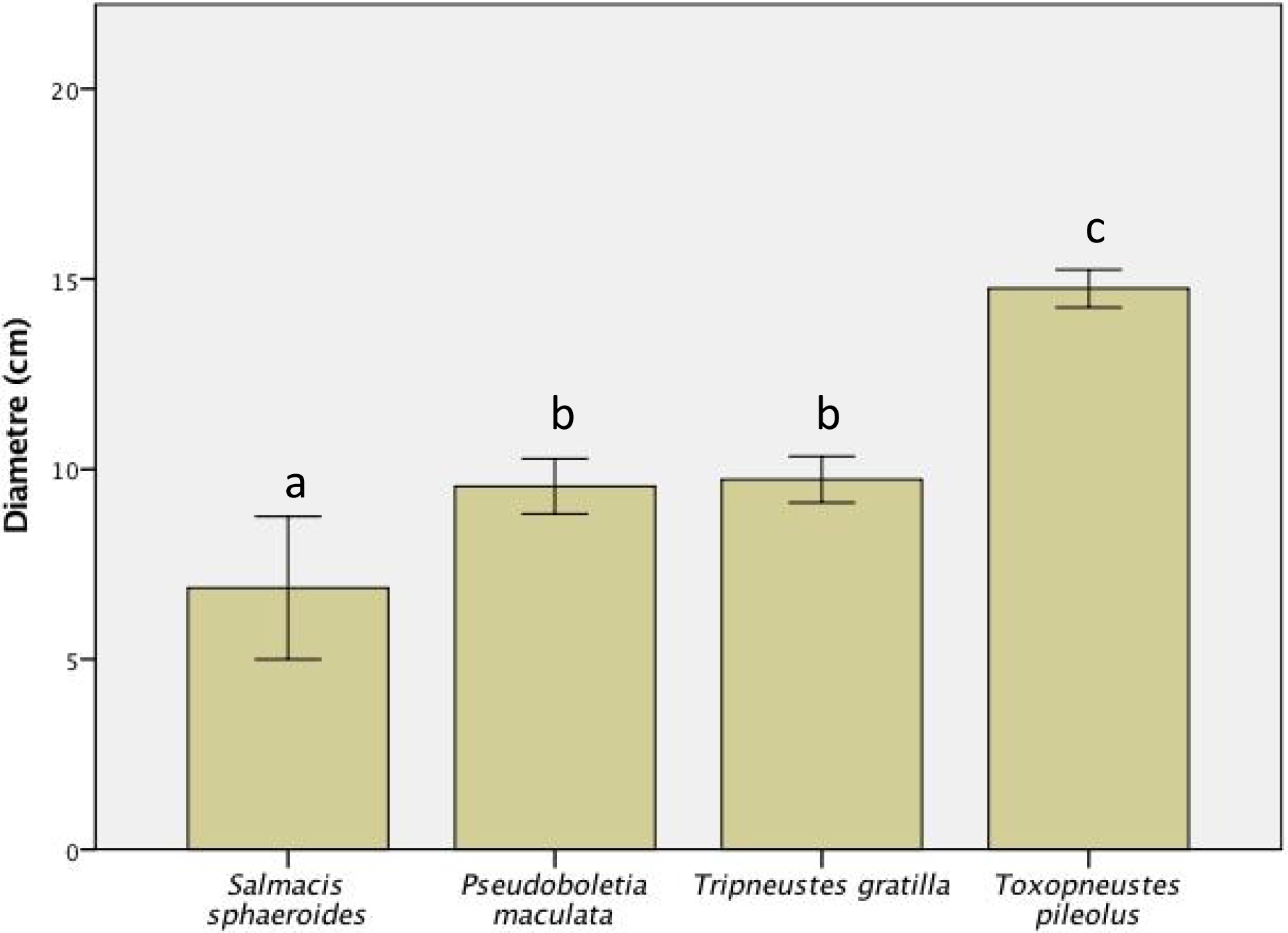
Diameter (mean ± SEM) of collector urchins. Means with different letters are significantly different (Tukeys HSD, p < 0.05).

**Figure 5:**
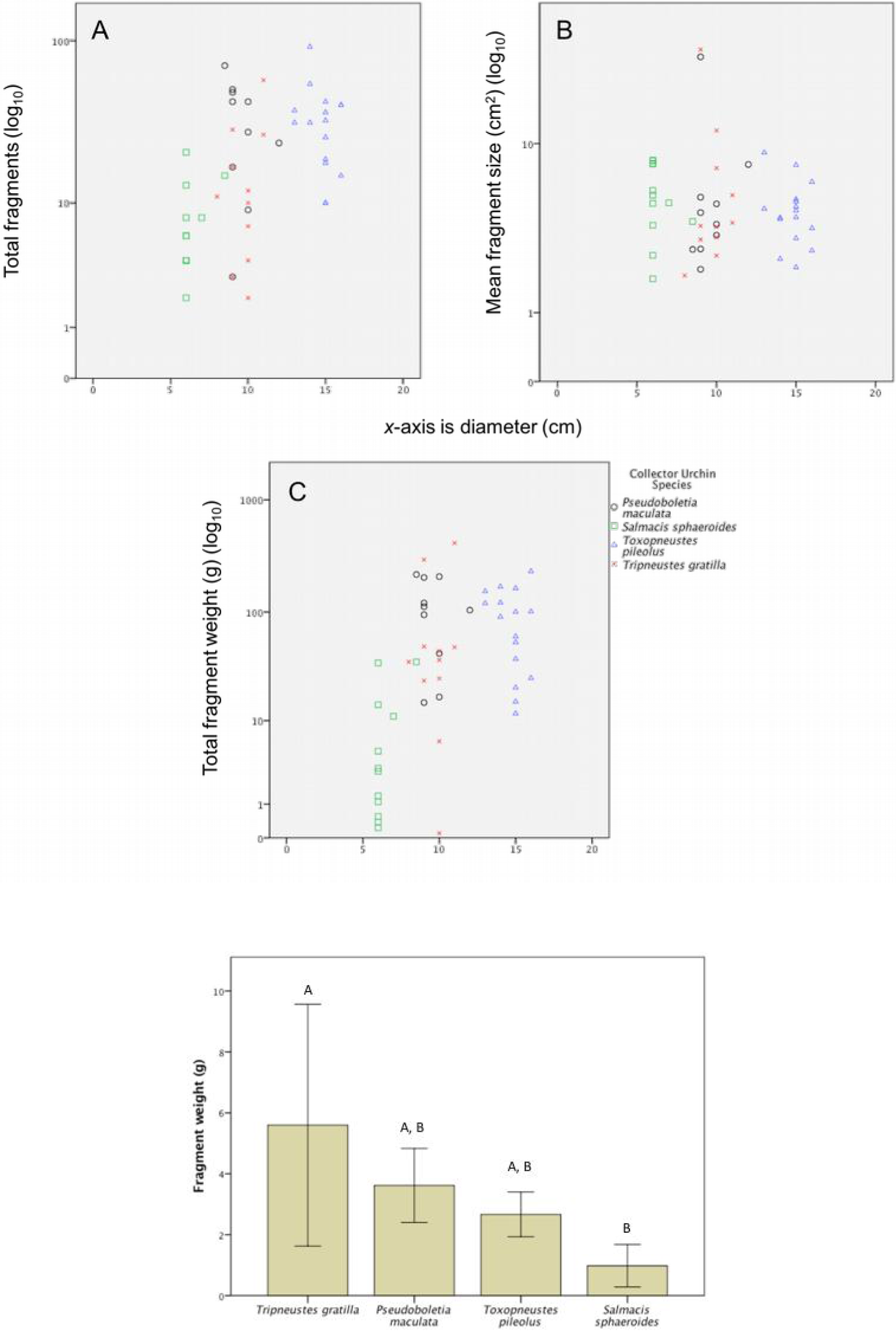
(A) Diameter v Total fragments; (B) Diameter v Average size of fragment (cm^2^); (C)Diameter v Total weight of fragments (g); *x*-axis is diameter (cm); *y*-axis is log_10_ scale. (D) Average fragment weight per species

Mean fragment weight (±SEM) in g per species was as follows: *Pseudoboletia maculata*: 3.62 ± 0.54, *Toxopneustes pileolus*: 2.67 ± 0.34, *Tripneustes gratilla*: 5.6 ± 1.78, and *Salmacis sphaeroides*: 0.98 ± 0.32. An analysis of variance (ANOVA) of the observed fragment weights yielded significant variation among urchin groups, F(3, 45) = 4.615, p = 0.007. A post hoc Tukey test, denoted by letters above error bars, shows significance among groups at p < .05. Mean difference in fragment weight of *S. sphaeroides* and *T. gratilla* was significant at the value of p = 0.004.

After we removed the debris coverage from the body of an urchin, it would quickly grab new debris with its tube feet and place it on top of its body. The speed of fragment movement onto the urchin was similar between different species (Fig.6A). In the footage we analyzed, *P. maculata* moved coral rubble the fastest, at 2.0 cm min^−1^, followed by *T. pileolus* at 1.91 cm min^−1^, and *T. gratila* at 0.99 cm min^−1^. The transport speed was not constant during the movement of a fragment, often with a period of fast transport being preceded and followed by slower movement (Fig. 6B). Hence, despite the divergent global speeds noted above, instantaneous transport speed (as seen in the slopes of the fragment positions in Fig. 6B) was relatively similar. Movements orienting the fragment also affected the speed of its movement towards the dorsal pole of the echinoid.

**Figure 6:**
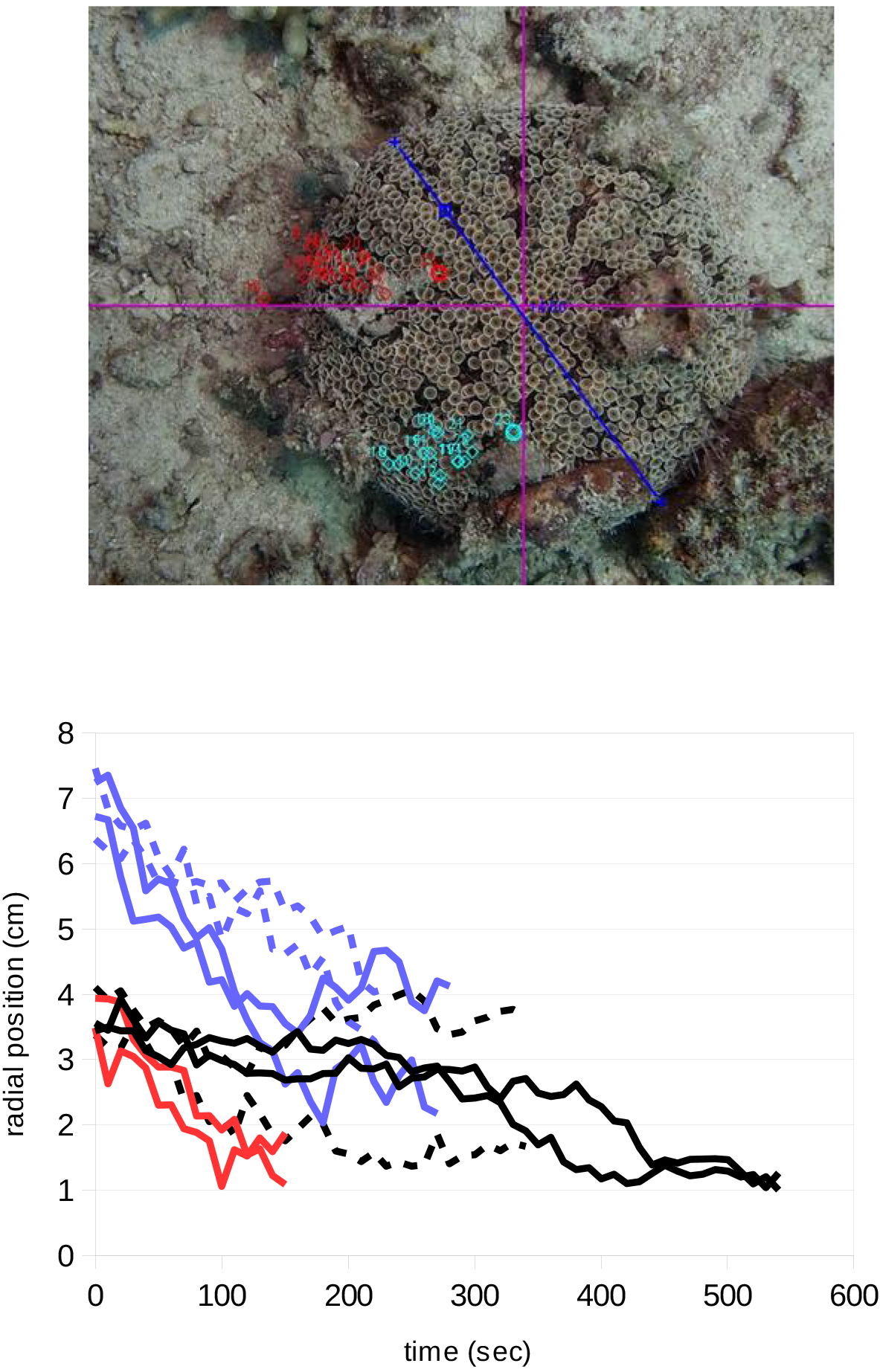
Speed of recovery of debris cover of urchins. A: Points indicating movement trajectory of the pole-most point of a coral rubble fragment on *Toxopneustes pileosus* (in the field). Note the scale bar in blue. Screen-shot from Tracker software. B: position of fragment relatives to the dorsal pole of the urchin. Species are color coded (P. *maculata*, red, *T. pileosus*, blue, *T. gratila*, black), continuous and broken lines indicate fragments on different animals. Footage recorded in tanks (*T. gratila* solid line, both *S. spheroides*) and in the field (all other recordings).

The covering behavior is jointly carried out by spines and tube feet. Initially the tube feet make searching rotational movements, until one of them contacts a piece of debris. The tube foot is soon joined by several other tube feet, which jointly pull the fragment to ward the urchin’s body (Fig. 7A,B). It is not obvious from the video observations if that is due to a directed recruitment of tube feet by the echinoid nervous system, or simply due to the proximity of neighboring tube feet to the detected fragment. Next, the tube feet pull the debris fragment closer to the echinoid. The fragment is most commonly transported up onto the body of the urchin by a combined pulling movement of the tube feet and a pushing movement of the spines. Upward-angled spines keep the fragment from slipping off, while the spines above the fragment part. The tube feet above the fragment are often seen pulling it towards the pole (anus) of the echinoid (Fig. 7C, D). The pedicallariae are not involved in the transport process. The fragments which reach the top of the urchin first seem to block the transport of further fragments, a process which repeats until the complete dorsal surface of the urchin is covered. We often observed that the urchins moved in one direction when stripped of covering material, while re-covering. This combined covering-walking behavior happened both in the field and in tanks.

**Figure 7:**
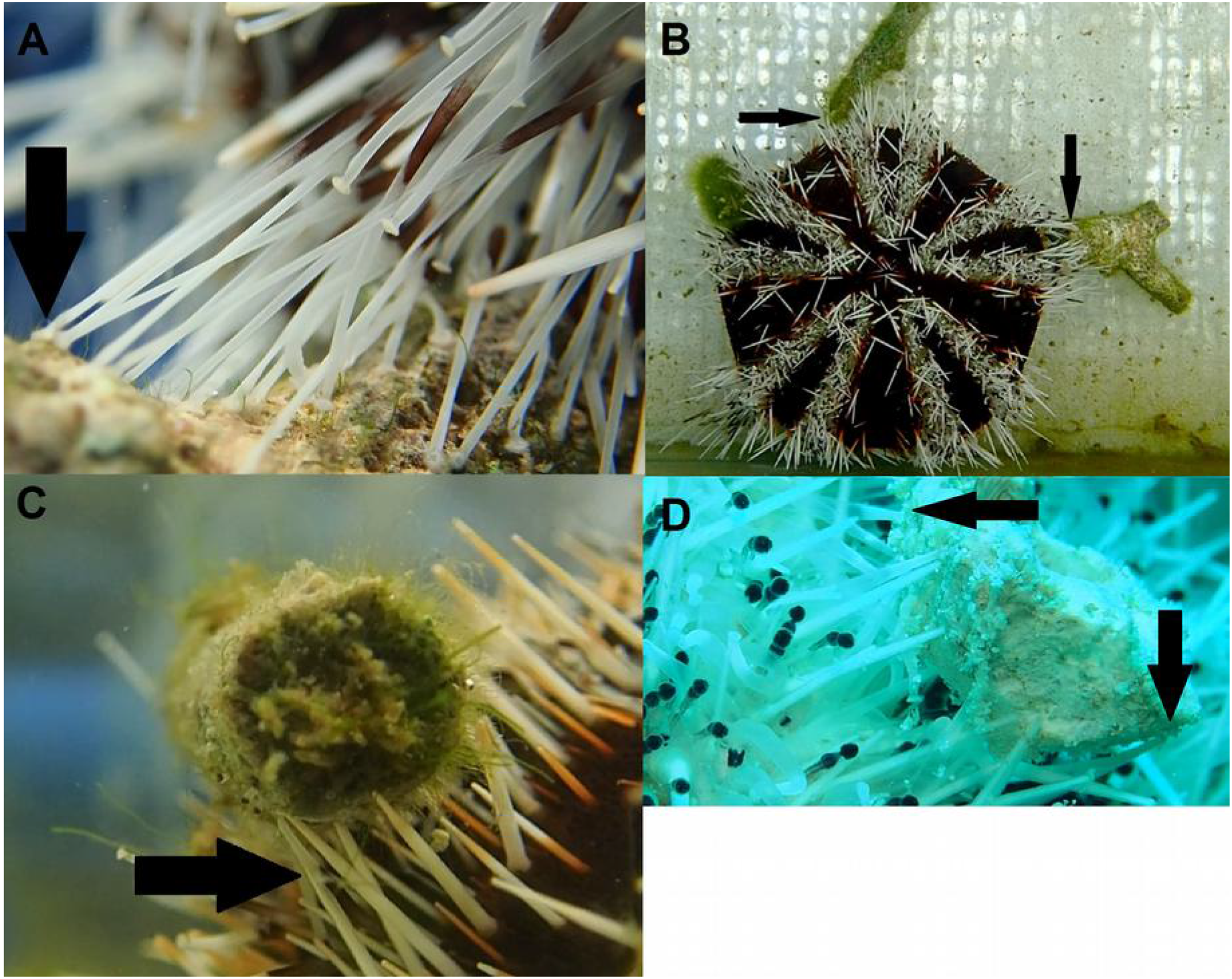
Spines and tube-feet are involved in the covering behavior in *Tripneustes gratila.* Screenshots of videos. A: several tube feet (one indicated by an arrow) jointly grab a coral rubble fragment in the initial phase of transport. B: two fragments held by tube feet (arrows). C: Spines pushing a fragment upwards. D: Spines pushing (downward arrow) and tube feet pulling (leftward arrow) a fragment. A, D are from footage recorded in the field, B, C in tanks.

Several anecdotal observations during our study are worth mentioning. Two of the urchins carried seemingly living tunicates on their backs when we found them. We found many urchins in areas with sand or coral rubble, interspersed by rocks not higher than 30 centimeters. Tunicates sit on the edges of many of these rocks, exactly in a position where a passing urchin would be in the position to impale them. Another observation we repeatedly made was that urchins stripped of debris would crawl underneath the protective spines of groups of nearby *Diadema* echinoids. Also, when the collector urchins replaced the debris covering their bodies after we removed it, they caused significant perturbation of the sand and rubble they walked on. The urchins were then often followed by wrasses (mostly *Choris batuensis, Oxychelinus sp., Diproctacanthus sp*.) and sand perches (*Parapercis sp*.), presumably hoping to gain food items from underneath the upturned substrate.

**Figure 8:**
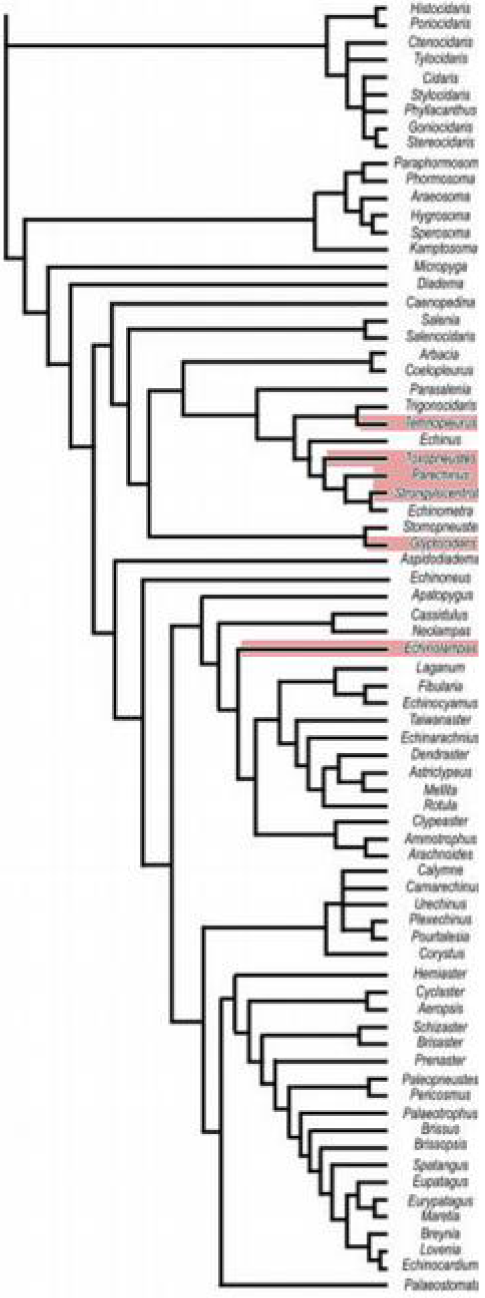
Phylogeny of covering behavior in echinoidea. Cladogram from (Kroh and Smith, 2010). Groups with species known to show covering behavior are highlighted in red. Marked, from top to bottom, are the Temnopleuridae, Toxopneustidae, Parechinidae, Strongylocentrotidae, Glyptocidaridae (regularia) and Echinolampadidae (iregularia). The branch point between regularia and irregularia echinoids lies about 200 ma in the past, the branch point between the Glyptocidaridae and the other regularia showing covering behavior occurred just slightly later.

## Discussion

Our study shows that the covering material of four species of collector urchins, *Pseudoboletia maculata, Toxopneutes pileolus, Salmacis sphaeroides* and *Tripneustes gratilla*, is distinctively species dependent. A previous study of the covering behavior of *Tripneustes ventricosus* and *Lytechinus variegatus* in the Caribbean found that *Tripneustes* prefers to cover itself with seagrass, even though it uses other material as well if little seagrass is available (Amato *et al.*, 2008). Our study comes to a similar conclusion, with 15% of the cover of the closely related *Tripneustes gratilla* being seagrass, compared to 0.5% and no seagrass cover for *Toxopneutes pileolus* and *Pseudoboletia maculata*, respectively. *Salmacis sphaeroides* had a higher preference for organic material with a total of 52%, 87% of this being seagrass alone. Our study hence confirms a species specific choice in covering material. The fact that the fragment size was similar for different urchin species of different sizes points to similar covering mechanisms in all these species. Video analysis of fragment transport agrees with this conclusion, and shows a covering mechanism involving both spines and tube feet.

We only found an increased debris coverage with depth in *Toxopneutes pileolus*, which confirms the purpose of the covering behavior as UV protection and weighting down to counter water movement seen in previous studies (Adams, 2001; Dumont *et al.*, 2007; Belleza, Samuel and Jr, 2012; Ziegenhorn, 2016). The limited depth range at which we sampled the other species might have contributed to the lack of depth dependence of coverage which we found.

### Echinoid covering behavior evolution

Covering behavior is phylogenetically widely occurring in echinoidea. The species compared here belong to the families of the Toxopneustidae (*Pseudoboletia maculata* and *Toxopneutes pileolus*), and Temnopleuroidae (*Tripneustes gratilla* and *Slamacis sphaeroides)*. Other echinoid families with members showing covering behavior are listed in table 1. These families span several groups of echinoids outside of the more primitive pencil urchins (cidaorids), and the long-spined diademadids (Fig. 7). Among the regularia (radially symmetric sea urchins), covering behavior is found in several groups of the Echinacea. A big cluster of collector urchins is found in the Camarodonta, including collectors in the families of the Parechinidae, Strongylocentrotidae, Temnopleuridae and Toxopneustidae. We found one mention of an irregular echinoid (asymmetric sea urchins, sand dollars and their relatives) with covering behavior, the deep sea urchin *Conolampas sigsbei*, belonging to the family of the Echinolampadidae in the Neognathostomata branch of the irregularia (Zhao *et al.*, 2013). We are aware that many irregularia bury themselves in the sediment. This behavior which likely shares mechanistic features with the covering behavior discussed here is nevertheless distinct in its ecological role, and we treat the two behaviors separately. The ancestors of these collector urchin families split around 200 ma ago in the lower Jurassic (Kroh and Smith, 2010). Since collecting behavior is not observed in the majority of echinoids, it presumably evolved convergently multiple times in the lineages mentioned here (For an extensive discussion of convergent evolution see (McGhee, 2011).

**Table 1:**
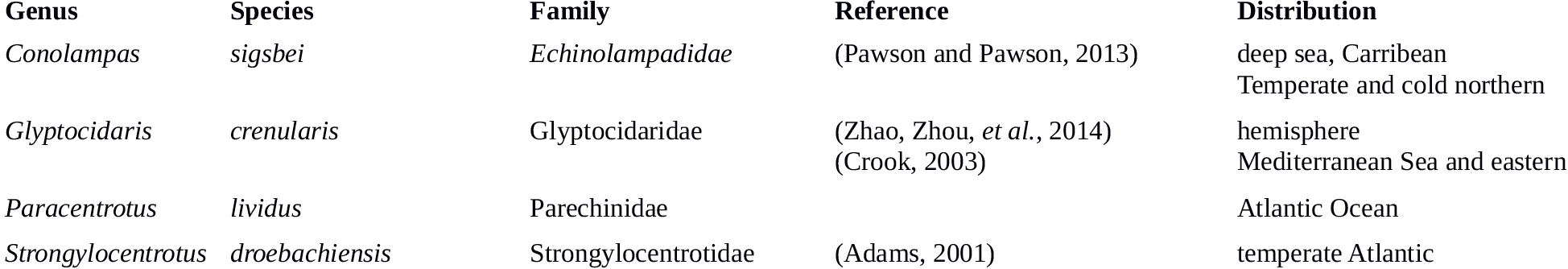
A list of echinoids showing covering behavior, assembled from the literature and from personal observations. We make no claim that this list is exhaustive. See also Fig. 7.

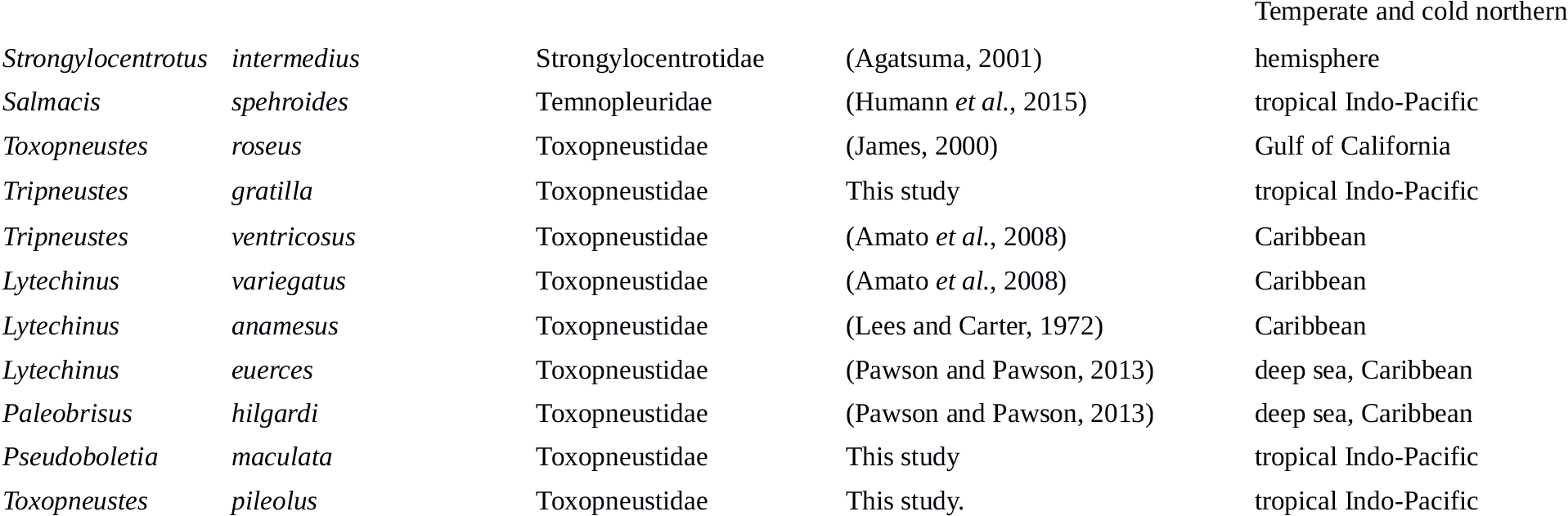

### Echinoid covering behavior as tool use

An intriguing question is if the covering behavior shown by urchins can be classified as animal tool use. Animal tool use is notoriously difficult to define, especially in borderline cases (Smith and Bentley-Condit, 2010). Following previous work by (Millikan and Bowman, 1967) and (Van Lawick-Goodall, 1971), (St Amant and Horton, 2008) expanded the definition by (Beck, 1980) and defined tool use as “… the exertion of control over a freely manipulable external object (the tool) with the goal of (1) altering the physical properties of another object, substance, surface or medium (the target, which may be the tool user or another organism) via a dynamic mechanical interaction, or (2) mediating the flow of information between the tool user and the environment or other organisms in the environment”. Collector urchins control freely manipulable marine debris of their choice with the goal of altering the physical properties of themselves (the tool user), for the sake of UV, predation and water surge protection. They also mediate the flow of information between the tool user and the environment or other organisms, by using the debris as camouflage.

The collecting behavior of urchins thus falls into two categories, as given in (Smith and Bentley-Condit, 2010), namely physical maintenance, the use of a tool to affect one’s appearance or body, and predator defense the use of a tool to defend oneself. Bentley-Condit & Smith cite the placement of anemones on their shells by hermit crabs as an example of the latter, which is quite similar to the echinoderm behavior discussed here.

A significant amount of reasoning an experimentation has been directed to the question how much reasoning and insight are behind animal tool use. Tool use by humans and presumably apes and crows derives from a mental model of the physical effects of the tool (reviewed in (Seed and Byrne, 2010)). Such an internal model however is not a condition for tool use, and Seed and Byrne reason that insect tool use (for example an ant lion flicking sand at prey) comes about without an internal representation of the tool as an extension of the animal’s body. Collector urchins most likely have no mental representation of the tools they are using or the intended consequences of the tool use, since they are lacking a centralized brain. Nevertheless, several observations indicate that echonoid covering behavior is tool use, as defined in the literature cited above:

- The use of *selected* pieces of debris as a cover by urchins (as shown in this study and in (Amato *et al.*, 2008; Ziegenhorn, 2016). By no means is the covering material simply debris which got stuck on the urchin, but it is there by choice of the animal, as a consequence of the coordinated action of tube-feet and spines.
- The *adjustment of debris cover according to need* (increased coverage in response to increased UV radiation, Dumont *et al.*, 2007, and the increased coverage at shallower depth, this study) also points to goal-directed tool-use behavior.
- The *difference in debris type between echinoid species* points toward a tool-use tailored to the needs and capabilities of the different urchin species.

We are of course by far not the first ones to observe and describe the covering behavior of urchins, but the conceptual connection between this behavior and tool use has, to our knowledge, not been made previously, besides a passing mention in (Mann and Patterson, 2013). We do believe that this behavior can be classified as tool use, adding the echinodermata as a fourth phylum with members capable of tool use next to the chordata (mainly primate mammals and crovid aves), arthropoda and molluska (primarily octopod cephalopods) (Smith and Bentley-Condit, 2010).

## Author Contributions

All authors collected data, G.B. conceived the idea, G.B. and K.M.S. analyzed the data and wrote the manuscript.

## Acknowledgments

We thank everyone at People and the Sea and at the Bolinao Marine Lab for their support, and Drs. Cecilia Conaco and Patrick Cabaitan helpful discussion of the manuscript.

